# A 3D *in vitro* cortical tissue model based on dense collagen to study the effects of gamma radiation on neuronal function

**DOI:** 10.1101/2023.06.07.544083

**Authors:** Neal M. Lojek, Victoria A. Williams, Andrew M. Rogers, Erno Sajo, Bryan J. Black, Chiara E. Ghezzi

## Abstract

Studies on gamma radiation-induced injury have long been focused on hematopoietic, gastrointestinal, and cardiovascular systems, yet little is known about the effects of gamma radiation on the function of human cortical tissue. The challenge in studying radiation-induced cortical injury is, in part, due to a lack of human tissue models and physiologically relevant readouts. Here, we have developed a physiologically relevant 3D collagen-based cortical tissue model (CTM) for studying the functional response of human iPSC derived neurons and astrocytes to a sub-lethal radiation exposure (5 Gy). We quantified cytotoxicity, DNA damage, morphology, and extracellular electrophysiology. We reported that 5 Gy exposure significantly increased cytotoxicity, DNA damage, and astrocyte reactivity while significantly decreased neurite length and neuronal network activity. Additionally, we found that clinically deployed radioprotectant amifostine ameliorated the DNA damage, cytotoxicity, and astrocyte reactivity. The CTM provides a critical experimental platform to understand cell-level mechanisms by which GR affects human cortical tissue and to screen prospective radioprotectant compounds.

## Introduction

Human exposure to high doses (>1 Gy) of gamma radiation (GR) primarily occurs in cancer-related radiation therapy. However, nuclear or radiological attacks and accidental exposures represent rare but critical modes of exposure, leading to acute to long-term injury or death. Understanding the biological mechanisms behind GR-induced injury is paramount for both medical and public health preparedness. After September 11, 2001, several research programs were established in the U.S. to ensure medical readiness for civilian casualties in the event of an ionizing radiation (IR)-related public health emergency (1). Specifically, research efforts have targeted improved understanding of radiation’s impact on systems throughout the body, and how to diagnose, mitigate, and treat resulting injuries (2–4). Most IR-induced injury studies have addressed one or more of the following systems/tissues: bone marrow, gastrointestinal tract, lungs, skin, vasculature, and kidneys (5–8). However, IR, including GR, is also associated with radiation-induced cognitive decline (RICD) (9); likely attributable to disruption of ‘healthy’ neuronal network function.

Longitudinal studies on survivors of the atomic bomb attacks on Hiroshima and Nagasaki, as well as the Chernobyl Nuclear Power Plant disaster, confirm direct deleterious effects on cognition, including acute-to-long-term memory loss and cognitive impairment (10, 11). Post-incident reports indicate survivors were primarily exposed to GR (12, 13). However, due to the nature of these incidents and the impracticalities of fundamental studies on human patients, little is currently known regarding the precise mechanisms leading to RICD (14). To date, mechanistic studies have largely been carried out in either animal models or 2D culture systems(14) (4). Of the few clinical and preclinical reports aiming to address underlying mechanisms, challenges regarding qualitative data collection and human disease relevancy of the model systems persist. Animal models to investigate mass casualty responses have relied on radiation survival colonies, comprised primarily of rodent, canine, or non-human primate models (1, 15, 16). In addition to important species-specific CNS cell/network differences (17), these critical cohort models rely on qualitative neurobehavioral assessments, are low-throughput, and are time and cost-intensive. Importantly, many of these studies simply fail to recapitulate the ionizing radiation (IR)-dependent neurocognitive decline (18, 19). More recently, *in vitro* CNS models have been used to study IR damage, in general (20, 21). However, these studies have primarily been carried out on 2D substrates, using tumorigenic cell lines or embryonic animal tissues (22), and have reported measurements of DNA double-strand break (DSB) and/or cell survivability only (20, 21, 23). It is important to note that (1) low-energy GR interacts with biological tissues in a depth-dependent (3D) manner, (2) tumorigenic and embryonic animal cells have limited pathophysiological relevance, and (3) DNA-damage and overall survivability are neither direct nor high-content measures of functional neuronal network activity.

In addition to a fundamental knowledge gap related to GR’s effect on cortical tissue, there is a clinical and military need to identify potential radioprotectant compounds. At high absorbed doses, acute radiation syndrome (ARS), is the clinical manifestation of IR-induced injury with onset within hours to days (24), and is divided into five sub-syndromes including hematopoietic, gastrointestinal, pulmonary, cutaneous, and neurovascular syndromes based on which organs have been primarily affected (25). The FDA currently approves four compounds for the treatment of hematopoietic-ARS and each of these approved pharmacological treatments are intended to mitigate the effects by stimulating myeloid progenitors within bone marrow (1, 26). To date, there are no FDA approved compounds for treatment of any non-hematopoietic-ARS sub-syndromes, including for symptoms classified as neurovascular-ARS (27). A number of pharmacological mitigations and cognitive therapeutic rehabilitation strategies have been studied clinically, with limited success (28). However, there remains no intervention approved for the treatment of central nervous system (CNS) disruption after GR exposure, including RICD. This is likely due to the lack of fundamental knowledge regarding the underlying biological and biochemical processes elicited by interaction between healthy human neuronal tissue and GR (29). In total, these points highlight a critical technology gap in the lack of 3D, physiologically relevant, quantitative, phenotypic experimental platforms for studying the biological response mechanism elicited by gamma-CNS tissue interactions and screening potential therapeutic compounds.

Here, we describe a novel 3D, human cell-based cortical tissue model (CTM) and gamma irradiation workflow that enables high-content phenotypic assessment of neuronal network function. The 3D *in vitro* CTM was initially seeded as proof of concept for initial evaluation with SH-SY5Y cells. Subsequently, CTM was seeded with a co-culture of human-derived induced pluripotent stem cells (iPSC) neurons and glia, resulting in native tissue architecture, composition, structural and mechanical features, and phenotypic traits. CTMs were cultured for as long at 64 days *in vitro* (DIV) and were irradiated with 5 Gy of GR; a dose relevant to acute survival of nuclear accidents or attacks (12, 30). We observed significant DNA damage, reduced neurite length, decreased firing and network activity in the CTM after GR treatment. Further, we effectively mitigated significant GR-induced injuries including increased DNA damage, and reduced neurite length by pre-treatment of CTM cultures with the known radioprotectant amifostine (250 µM). The described CTM represents a powerful experimental tool for understanding acute and chronic effects of gamma exposure on molecular, cellular, and neuronal network-level function as well as for evaluating the preliminary efficacy of potential therapeutic compounds. While the current work focuses on GR exposure, this tool is extensible to studies of mixed IR exposures relevant to both clinical tumor radiotherapies as well as studying the established and prospective neuronal effects of cosmic galactic radiation (31–34).

## Results

### CTM morphological and mechanical properties

Dense collagen CTMs were fabricated by plastic compression, that resulted in 3D structures with physiologically relevant protein content, while cells were incapsulated at point of production (Figure 1). To characterize morphological and mechanical properties of CTMs (Figure 2A), acellular samples were imaged using scanning electron microscopy (SEM). We observed suprastructural (quarter-staggered) organization of collagen fibrils, as expected for this biofabrication method (35, 36) (Figure 2B). Unconfined compression testing resulted in a maximum stress value of 50.88 kPa at strain of 80% (Figure 2C). The mean compressive modulus was measured to be 30.00 ± 8.1 kPa, which is in the range of native soft tissue (37, 38) (Figure 2D). Importantly, we did not observe substantial structural changes in diameter or thickness due to cell-mediated remodeling or swelling after 64 days of culture (Figure 2A). In addition, confocal laser scanning microscopy analysis of SH-SY5Y cells, positively stained with DAPI and β III tubulin, demonstrated long term viability up to day in vitro (DIV) 64 with extension of neuronal processes (Figure 2E).

**Figure 1.**
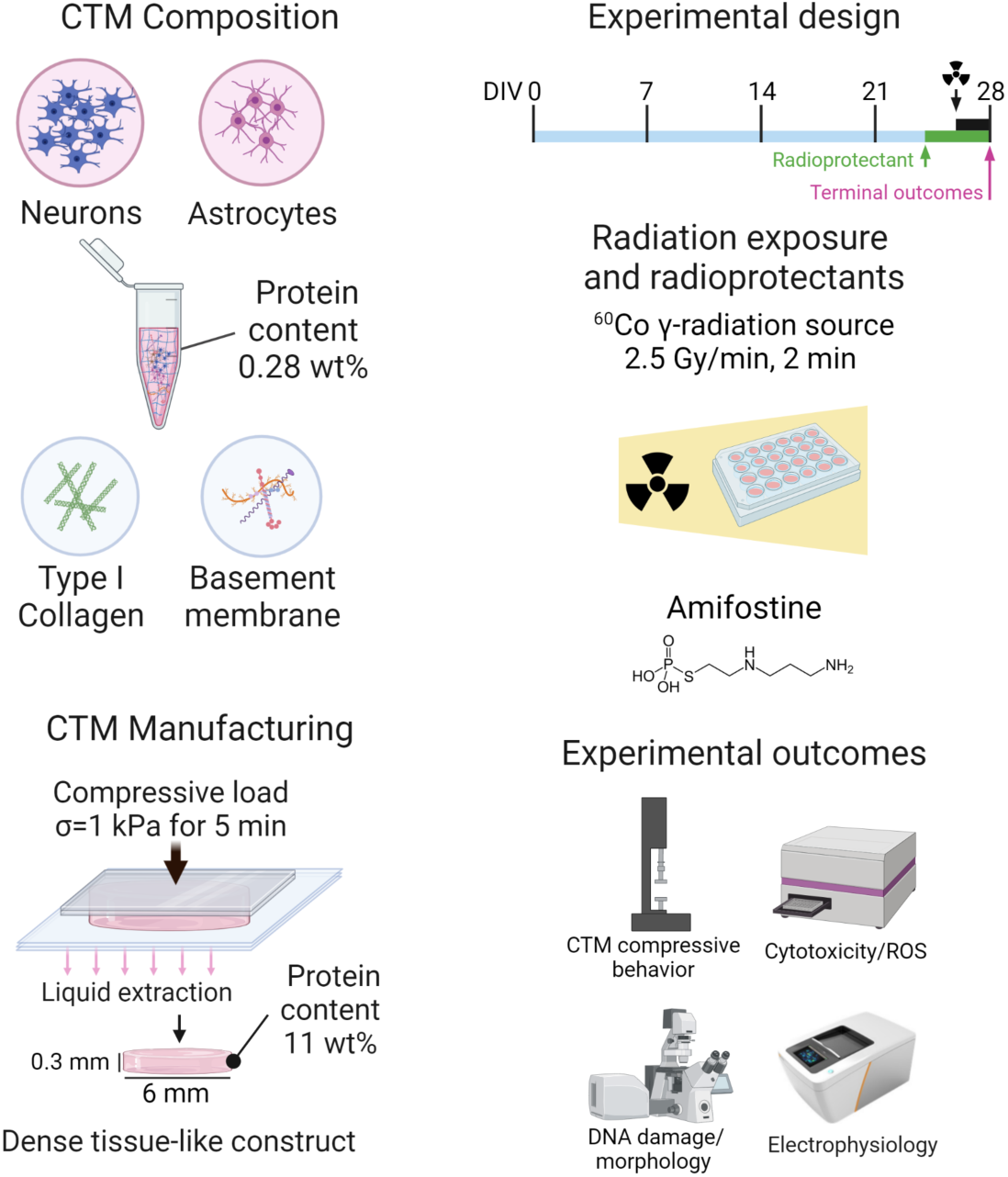
CTM preparation and experimental workflow. Cortical tissue models (CTMs) were generated by combining cells with rat tail type I collagen and basement membrane proteins. Upon gelation, highly hydrated collagen-based hydrogels underwent plastic compression to remove excess water and increase the protein content, resulting in a dense collagen-based CTM. After 26 days in culture, CTMs were irradiated with a Co^60^ GR source at 2.5 Gy/min for 2 min to simulate a 5 Gy dose to cortical tissue. Cultures were treated before radiation exposure with the commercially available radioprotectant (RP) amifostine by addition to culture media. CTMs were morphologically and mechanically characterized. The effect of GR on CTM function was quantified via cytotoxicity, DNA damage, cellular morphology, and electrophysiological behavior.

**Figure 2.**
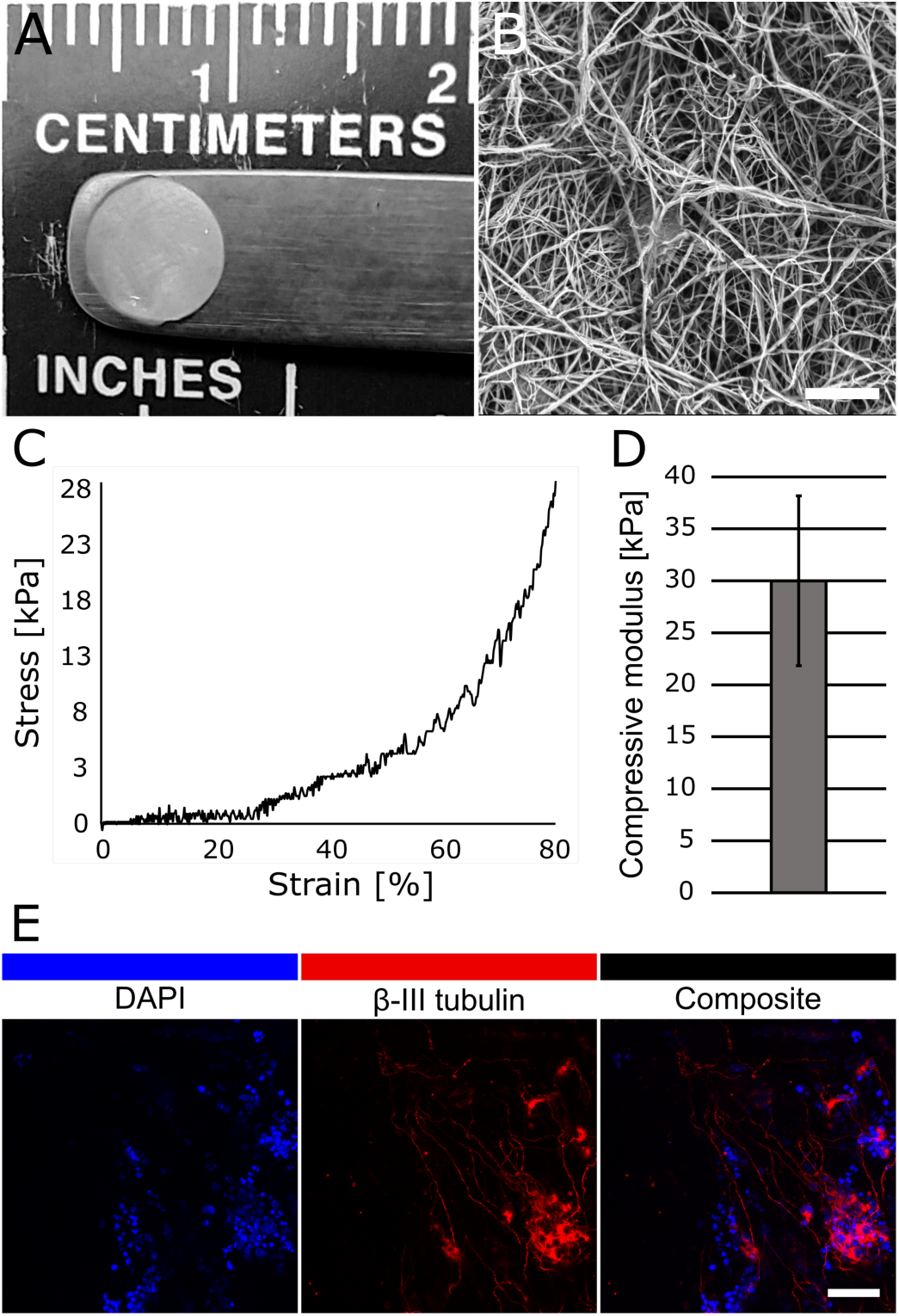
CTM morphological and mechanical characterizations. A. Macroscopic image of SH-SY5Y seeded CTM cultured to DIV 64. B. SEM analysis demonstrating suprastructural (quarter-staggered) organization of collagen fibrils. Scale bar represents 2 µm. C. Representative stress/strain curve of collagen-based constructs undergoing unconfined compression with compressive modulus in the range of native soft tissues (D). E. Immunocytochemistry staining for DAPI and β III tubulin in SH-SY5Y seeded CTM cultured to DIV 64. Scale bar represents 50 μm.

### GR-induced DNA damage and cytotoxicity

To simulate the effects of GR on human cortical tissue, we exposed SH-SY5Y-seeded CTMs to 2.5 Gy/min over 2 min from a ^60^Co source (UML Radiation Facility) for a total dose of 5 Gy. The primary mode by which IR sources, including GR, are known to induce cytotoxicity is through the induction of DNA damage. DSBs are recognized as the most deleterious form of DNA damage and are highly cytotoxic, resulting in partial gene deletion and/or translocation events (39, 40). To quantify DNA DSBs following GR treatment, we used ICC to evaluate the abundance of phosphorylated histone H2AX (γH2AX) and 53BP1, canonical markers of DNA DSBs, at 30 min post-treatment. Figure 3A shows representative images of ICC staining of CTMs seeded with SH-SY5Y for both γH2AX and 53BP1 DSB markers. Quantification of γH2AX and 53BP1 nuclei object intensity showed significantly greater intensity (4.29 ± 2.75 and 49.45 ± 23.92) in gamma treated CTMs than untreated controls (4.006 ±1.99 and 45.41 ± 23.19) (n=3, ROI = 5 per sample, 832 nuclei in irradiated sample and 212 nuclei in untreated sample) (Figure 3B). Levels of cytotoxicity in SH-SY5Y-seeded CTMs were quantified by measuring lactate dehydrogenase (LDH) release in culture medium at 24 h post-treatment (Figure 3C) and found to be significantly increased in treated samples.

**Figure 3.**
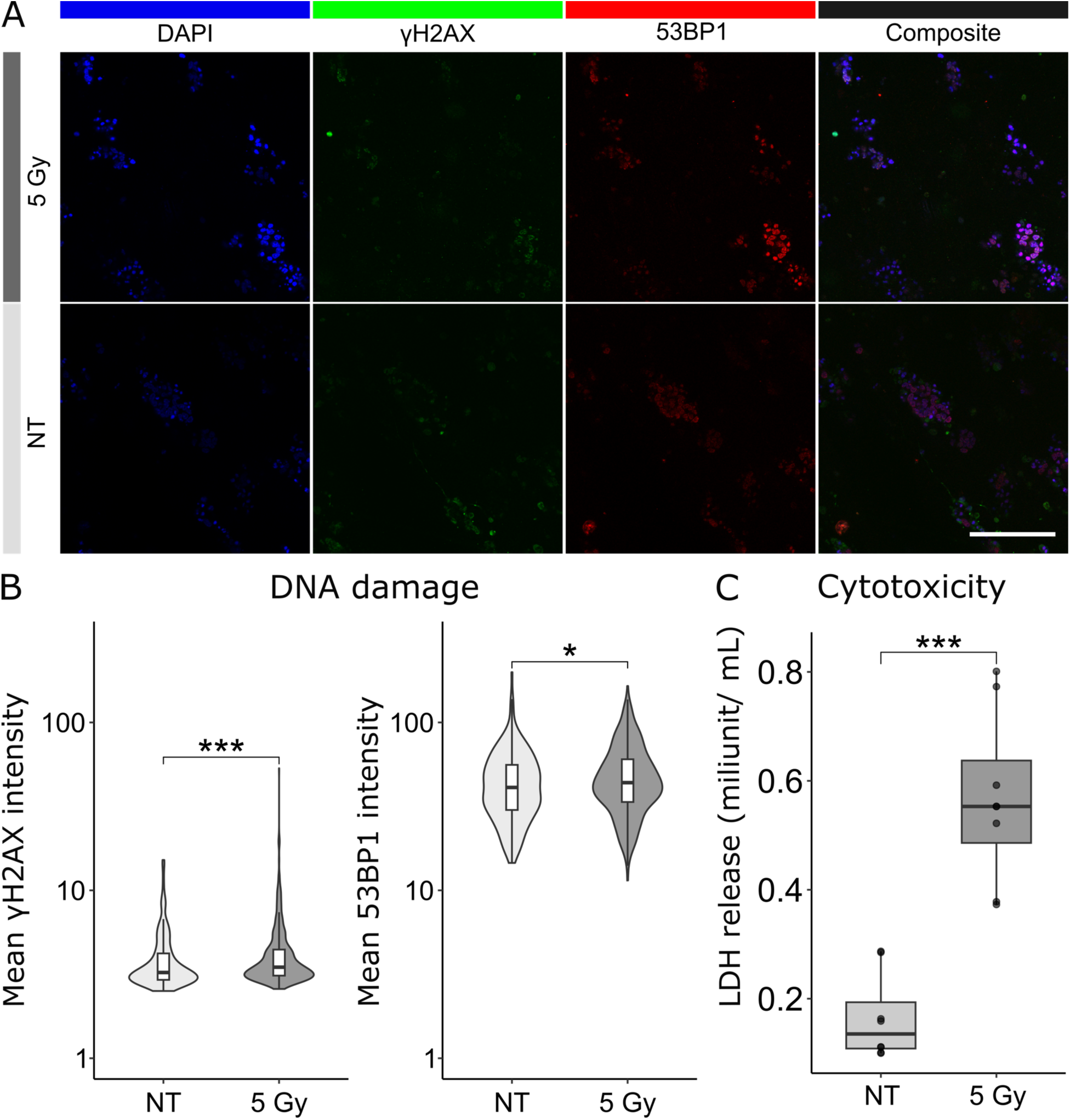
Biological response of SH-SY5Y -seeded CTM to GR. A. Representative maximum intensity projections of CLSM analysis of CTMs seeded with SH-SY5Y cells stained for DAPI (blue), γH2AX (green), and 53BP1 (red) at 30 minutes post treatment with 5 Gy GR. Scale bar represents 100 µm. B. Quantification of DNA damage markers γH2AX and 53BP1 mean intensity per nuclei at 30 min post GR treatment. Significant effect of radiation treatment on DNA damage marker expression in CTMs for both γH2AX (p = 0.00021) and 53BP1 (p = 0.018) nuclear intensities. C. LDH release from CTMs at 24 hours after treatment (p = 0.00093).

Patient-derived somatic cells can be converted into hiPSCs, capable of differentiating into a variety of cell types *in vitro*. hiPSCs have been introduced as a potential alternative for human- or animal-based therapeutic screening given their manufacturability, scalability, and ability to retain genetic characteristics of the host (41). CTMs seeded with hiPSC derived astrocytes and glutamatergic neurons were co-cultured for at least 26 days prior to 5 Gy of gamma irradiation. These cultures were likewise stained for DNA damage markers γH2AX and 53BP1 30 min post-treatment (Figure 4A). Quantification of DNA damage markers γH2AX and 53BP1 showed a significant increase in the intensity of both markers in the gamma irradiated (8.98 ± 8.61, 22.80 ± 13.56) versus the untreated CTMs (6.11 ± 5.61, 16.42 ± 11.13) (n = 5, ROI per sample = 7, NT = 1678 nuclei and 5 Gy = 2177 nuclei) (Figure 4B).

**Figure 4.**
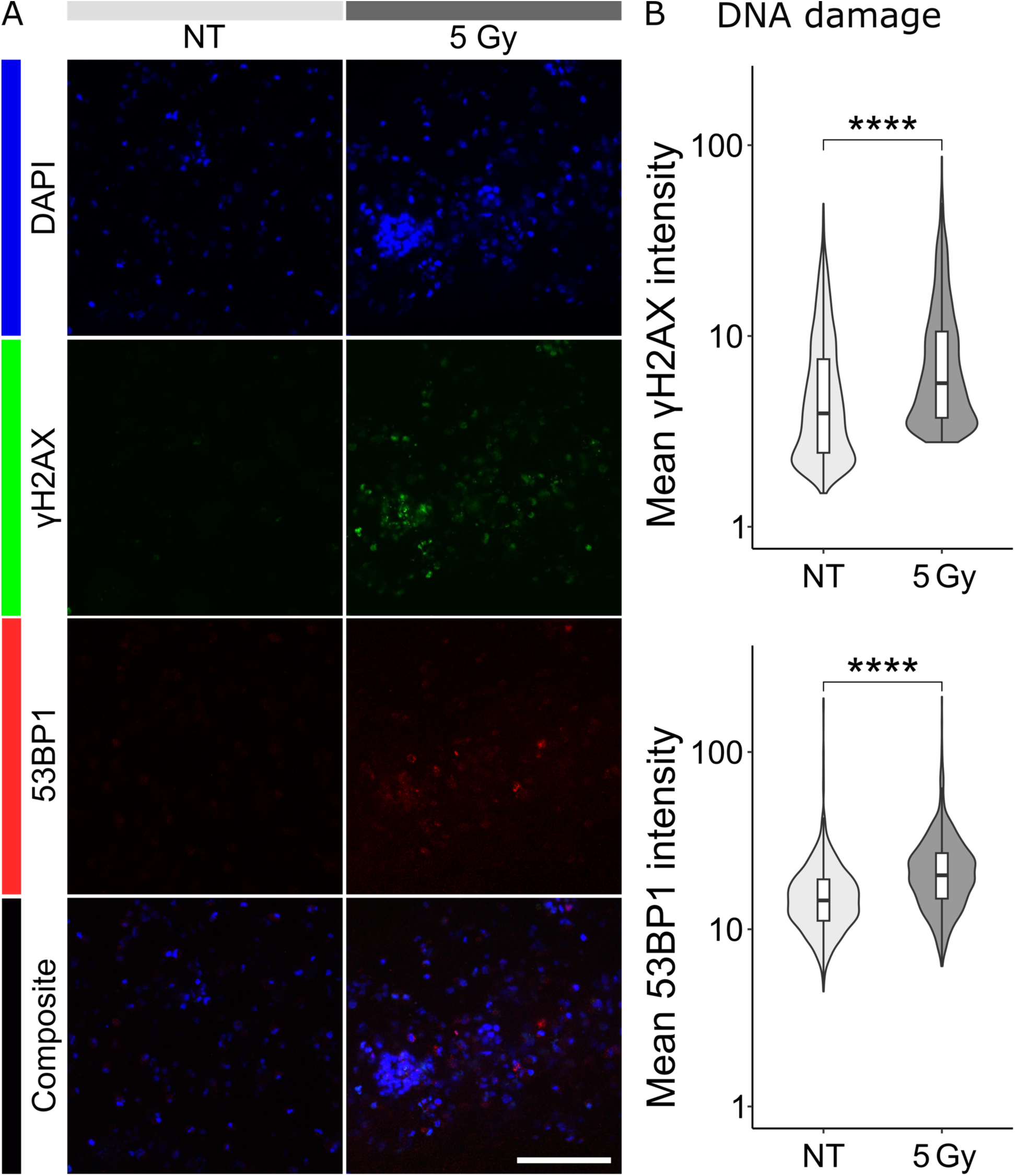
Biological response of hiPSC neuron and astrocyte-seeded CTMs to GR. A. Representative maximum intensity projections of CLSM analysis of CTMs seeded with hiPSC derived neurons and astrocytes stained for DAPI (blue), γH2AX (green), and 53BP1 (red) at 30 min post-treatment with 5 Gy GR. Scale bar represents 50 µm. B. Quantification of DNA damage markers γH2AX and 53BP1 mean intensity per nuclei at 30 min post GR treatment. Significant effect of radiation treatment on DNA damage marker expression in CTMs for both γH2AX (p = 2.22E-16) and 53BP1 (p = 2.22E-16).

### GR-induced changes in hiPSC morphology and reactivity

A further assessment of CTM morphological changes was made by ICC staining for the neuronal marker NeuN and the astrocyte markers GFAP (Figure 5A). A visible decrease in both neuronal and astrocytic processes is evident in the gamma irradiated CTMs. Quantification of neuronal processes length in NeuN revealed a significant decreased process length in irradiated (24.18 ± 16.39 μm) versus untreated CTMs (176.72 ± 53.92 μm) (irradiated samples = 20 measurements, untreated samples = 30 measurements) (Figure 5B). In addition, 91.52% cells are confirmed positive for the canonical neuronal marker TUJ1 (Figure 2E). To assess astrocyte reactivity, immunolabeled GFAP intensity was quantified. Astrocytes in GR irradiated CTMs exhibited significantly higher GFAP intensity than those in untreated controls (41.45 ± 2.56 versus 27.47 ± 2.09, p = 0.007).

**Figure 5.**
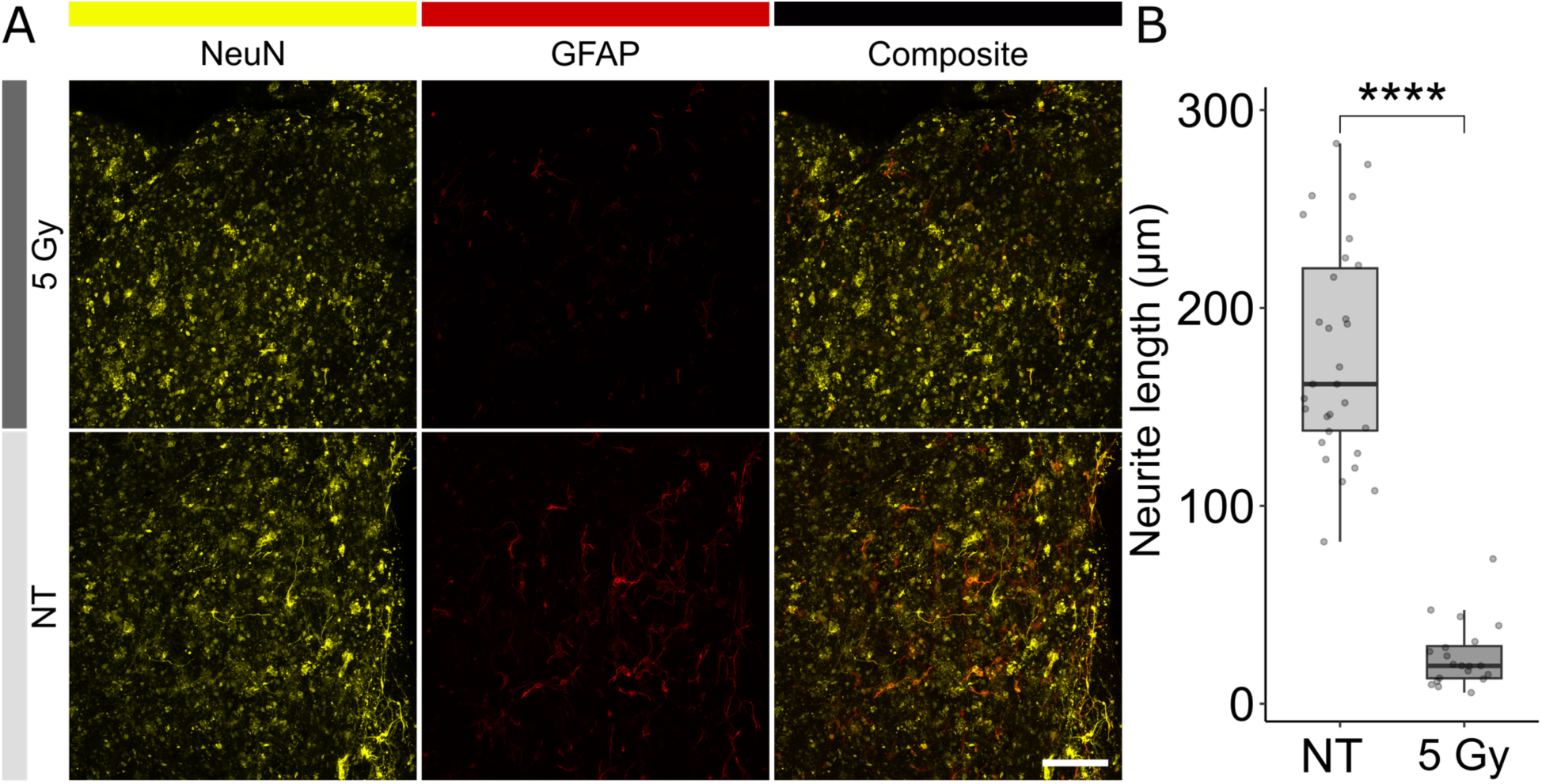
Morphological changes of hiPSC neuron and astrocyte-seeded CTMs to GR. A. Representative images of maximum intensity projections of CLSM analysis of CTMs stained for NeuN and GFAP. B. Significant effect of GR treatment in neurite length quantification (p = 4.4E-14) in CTMs relative to untreated samples at 24 h after treatment. Scale bars represent 50 µm.

### GR-induced changes in electrophysiological activity of 2D and 3D hiPSC co-cultures (CTMs)

To characterize the phenotypic activity before and after exposure, we cultured and irradiated both 2D and 3D co-cultures (CTMs) of hiPSC glutamatergic neurons and astrocytes. 2D co-cultures were seeded directly onto pre-treated multi-well MEA plates, while CTMs were transferred to a multi-well MEA plate for recording. Voltage versus time recordings were performed for both seeding conditions every alternate day for at least 26 days. 2D co-cultures began exhibiting extracellular action potentials (EAPs) on DIV 7 and exhibited a stable baseline of both mean firing rate (MFR) and network bursting rate (NBR) by DIV 27 (Figure 6C). While we did not observe single unit extracellular action potential (EAP) recordings in 3D-seeded wells, we did observe larger amplitude deflections consistent with multi-unit activity (MUA, Figure 6B) (42). MUA was responsive to typical agonists and antagonists of neural activity (Figure 6E), confirming biological sourcing of recorded activity. Subsequent recordings were carried out 1, 24, and 48 h following GR irradiation. We observed no significant differences in the rate of threshold crossings between irradiated and untreated CTMs (-12.51 ± 44.60 DoS, p = 0.93). However, we did observe a significant decrease in both MFR (3.75 ± 1.26 versus 2.96 ± 1.34, p = 0.04) and NBR (-53.85 ± 16.15 DoS, p = 0.04) in 2D co-cultures 24 h following irradiation.

**Figure 6.**
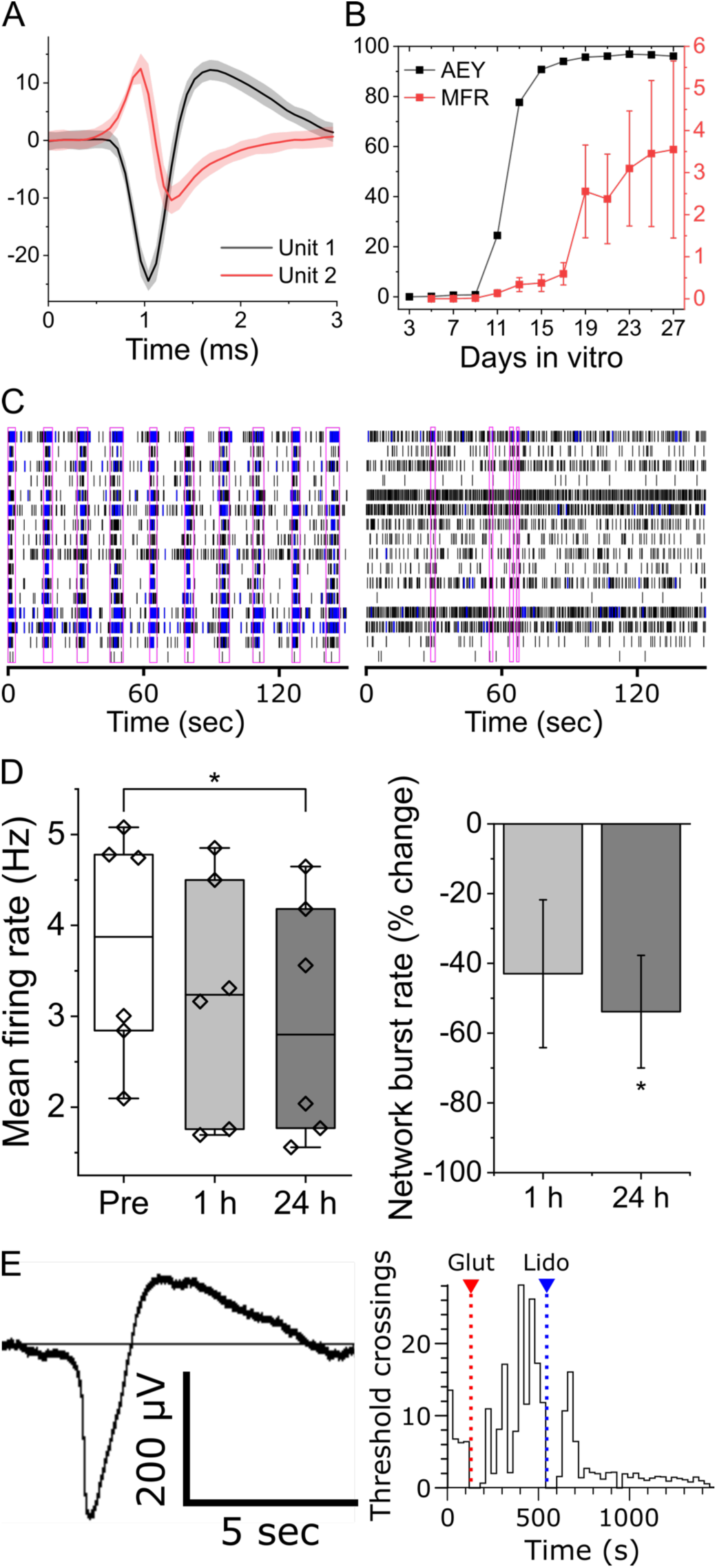
Electrophysiological activity changes of hiPSC neuron and astrocyte-seeded CTMs to GR. A. Representative voltage versus time traces for EAPs recorded from 2D co-cultures. Vertical and horizontal axes represent amplitude (µV) and time (ms), respectively. B. Time-progression and stability of 2D co-cultures in terms of AEY and MFR (left and right axis, respectively). C. Representative raster plots before (left) and 24 h (right) following GR exposure. Synchronized network bursts highlighted by pink rectangles. D. Difference-over-sum (DoS) normalized changes in MFR (left) and NBR (right) in 2D co-cultures at 1- and 24-hour following GR-exposure. E. Representative raw trace (left) of voltage versus time exhibiting MUA activity recorded from 3D CTMs. Pharmacological responsiveness (right) of 3D CTMs to glutamate (10 μM) and sodium channel antagonist lidocaine (100 μM).

### Mitigating GR-induced phenotypes in CTM with amifostine

To evaluate the dynamic response of CTM to radioprotectant drugs in radiation mitigation, we treated both hiPSC CTMs and 2D CTM co-cultures with 250 μM amifostine one hour prior to irradiation. CTMs were stained for the DNA damage markers γH2AX and 53BP1 at 30 min following irradiation (Figure 7A and B, respectively). Both γH2AX and 53BP1 associated intensities were measured for non-irradiated-vehicle (6.527 ± 11.13, 36.23 ± 20.79), non-irradiated-amifostine (9.32 ± 15.38, 35.32 ± 22.41), irradiated-vehicle (9.09 ± 14.90, 43.97 ± 32.06), and irradiated-amifostine conditions (6.62 ± 9.42, 33.84 ± 21.59) (n = 3 samples per condition, ROI per sample = 5, number of nuclei per condition: non-irradiated-no-radioprotectant = 1232, non-irradiated-amifostine = 1341, irradiated-no radioprotectant = 1439, irradiated-amifostine = 1466). 2D cultures and CTMs were stained for the neuronal marker NeuN (Figure 7C). We quantified neurite length in 2D non-irradiated vehicle treated cultures (391.63 ± 216.47 μm), non-irradiated-amifostine treated cultures (546.88 ± 166.06 μm) irradiated-vehicle treated cultures (330.77 ± 162.39 μm), and irradiated amifostine treated cultures (415.13 ± 184.38 μm). Amifostine had a significant positive effect on neurite length in irradiated 2D co-cultures (Figure 7D). Additionally, CTMs were stained for NeuN and neurites length were quantified in non-irradiated-vehicle treated CTMs (163.37± 84.06 μm), non-irradiated-amifostine treated CTMs (203.83 ± 59.56 μm), irradiated-vehicle treated CTMs (113.34 ± 63.85 μm), and irradiated amifostine treated CTMs (240.31 ± 122.60 μm). Amifostine had a significant positive effect on neurite length in irradiated CTMs (Figure 7E).

**Figure 7.**
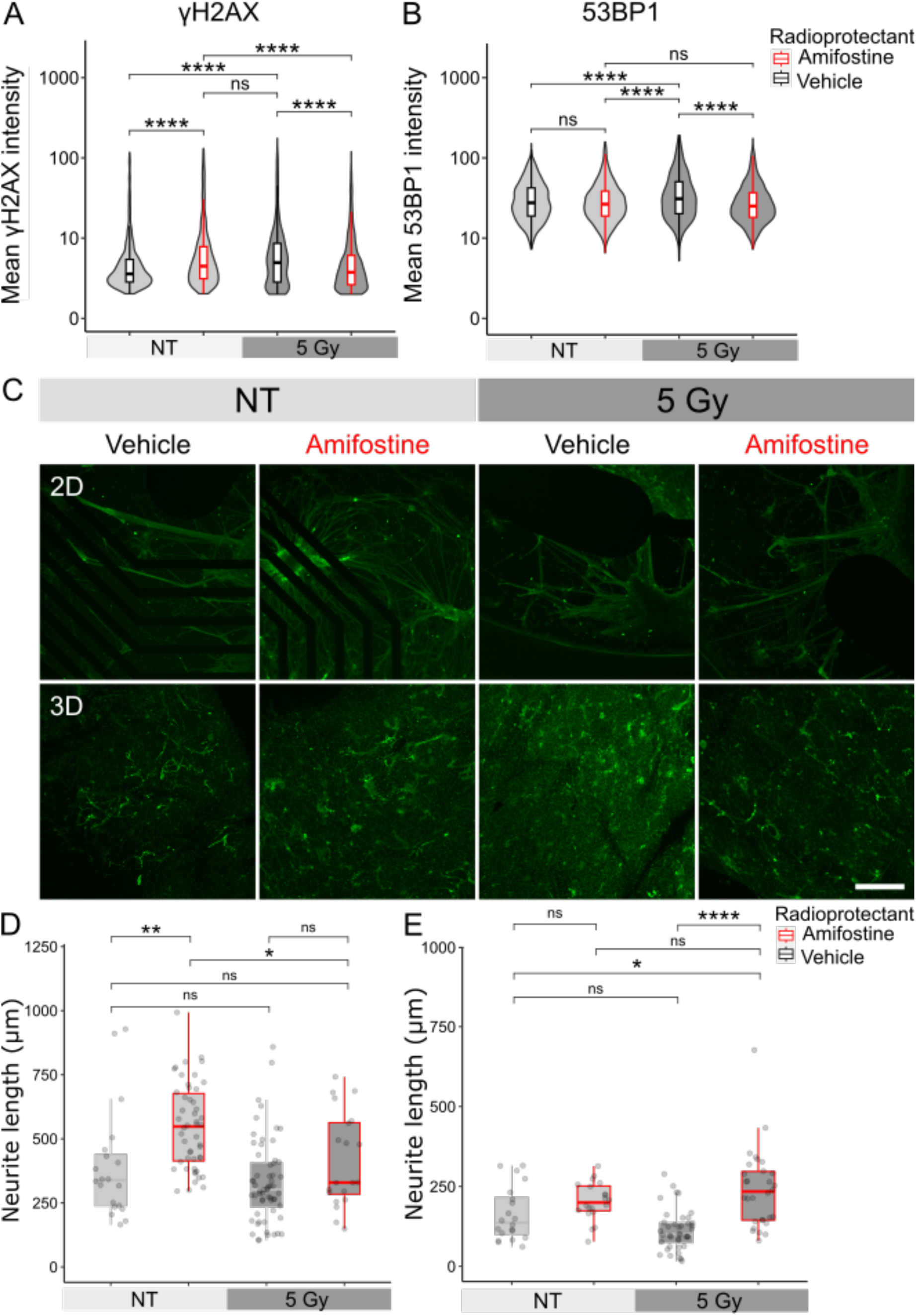
Biological response of hiPSC neuron and astrocyte-seeded CTMs to radioprotectant amifostine upon GR. Quantification of DNA damage markers γH2AX (A) and 53BP1 (B) mean intensity per nuclei at 30 min post GR treatment with and without radioprotectant. Significant effect of radiation treatment on DNA damage marker expression in CTMs for both γH2AX (p = 2.22E-16) and 53BP1 (p = 2.22E-16) and significant effect of amifostine on DNA damage marker expression in CTMs for both γH2AX (p = 5.055e-14) and 53BP1 (p = 7.51e-18). C. Representative images of maximum intensity projections of CLSM analysis of CTMs stained for astrocytic marker GFAP in gamma irradiated or untreated CTMs treated with the radioprotectant amifostine or vehicle. Scale bar represents 50 µm. Quantification of neurite length in 2D (D) and CTMs (E). Significant effect of amifostine in neurite length quantification (p = 8.191e-7) in CTMs relative to vehicle control upon RG exposure.

## Discussion

Despite the clear clinical, military, and commercial need to better understand the mechanisms of IR-cortical tissue interaction as well as to identify effective radioprotectant drugs, an *in vitro* CTM that recapitulates the structural, mechanical, and functional profile of the human cortex has yet to be established. Here, we report the development of a dense type-I collagen-based CTM that incorporates human basement membrane proteins and supports the growth, differentiation, and function of SH-5Y5 cells as well as human iPSC neurons and astrocytes. The optimized biofabrication technique (35, 36) produces cellular CTMs within minutes, which may be preferable to the days or weeks required by highly variable cell-mediated remodeling techniques (43). Although type-I collagen is not present in the brain, it is a widely used and well-characterized fibrillar protein that provides a biocompatible, tunable, structurally stable 3D environment, as biomanufacturing platform for CTM (35, 36). In the current manuscript, SEM and unconfined compression testing were used to evaluate the structural and mechanical properties of the CTM. Current fabrication strategy results in quarter-staggered suprastructural organization, similar to that previously reported (35), and a compressive modulus of 30.00 ± 8.1 kPa (Figure 2B). While this modulus is within the soft-tissue range (37, 38), it is slightly higher than that of native brain tissue (approximately ∼0.3kPa to ∼6 kPa)(38) . Future iterations of the CTM may include functional natural spacers materials, such as hyaluronic acid, to further tune the bulk modulus as well as allow tuning of the hydrophilicity and ECM protein distribution. While porosity and permeability were not explicitly tested, CTMs supported homogeneous cell distributions following seeding as well as the long-term viability of both cell types (at least 62 days in the case of SH-5Y5 cells). Importantly, but expectedly, we observed CTM structural stability over time, with limited-to-no cell-mediated remodeling.

To validate our model’s response to IR and its response to radioprotectant drugs, CTMs were seeded with either SH-5Y5 (as a proof-of-concept) or iPSC neurons and astrocytes and then exposed to a single dose (estimated 5 Gy) of GR in the presence or absence of the radioprotectant amifostine. Standard DNA DSB markers γ-H2AX and 53bp1 were immunolabeled and quantified 30 min following irradiation, to capture radiation-induced changes, as previously reported (44, 45). We observed significant increases in both γH2AX and 53bp1 in gamma-irradiated CTMs (Figures 3B) and, importantly, a baseline expression level of both markers in the case of amifostine pre-treatment, suggesting a protective role against DNA damage (Figure 7A and 7B). Upregulation and co-localization of γH2AX and 53bp1 are widely used markers for DSBs (46) in studies of general oxidative stress (47) as well as radiation-induced injury (48), as for sparsely ionizing GR-ROS generation and subsequent stresses are thought to be the primary mechanism of radiation-induced injury (49). Amifostine is a cytoprotective drug developed by the Antiradiation Drug Development Program (ADDP) of the US Army Medical Research and Development Command as a radioprotective compound. While there are no radioprotectant (or ‘radiation countermeasure’) drugs approved for clinical use by the FDA, amifostine has been approved by the ADDP and the FDA for non-clinical use as a radiation countermeasure (50). Amifostine’s primary mechanism of action is the scavenging of free radicals. However, its metabolite is also thought to provide radioprotection through rapid oxygen consumption and the subsequent intracellular anoxia (51).

To date, histological and *in vitro* studies of IR-induced neuronal/cortical damage have largely been limited to DNA DSB and/or cytotoxicity measurements (e.g., LDH assays). To further validate our CTM and expand the functionally relevant biological readouts that may be associated with radiation-induced injury, we quantified neurite length and astrocyte reactivity using ICC, and functional network activity using microelectrode arrays (MEAs). To the best of our knowledge, these are the first demonstrations of these readouts from an IR-irradiated CTM.

To evaluate morphological changes in our neuronal network (i.e., neurite length), we immunolabeled NeuN and manually traced and measured prominent neurites in both 2D and 3D CTMs. In this study, we observed significant reductions in neurite length following gamma irradiation (Figure 5B) in both 2D cultures and 3D CTMs. Additionally, we observed that neurite lengths were preserved in CTMs pre-treated with neuroprotectant amifostine (Figure 8 C, D, and E). Oxidative stress is widely known to induce neurite degeneration, both *in vivo* and *in vitro* (52, 53). Moreover, neurite degeneration results in cortical network dysfunction as synaptic connections are pruned (54). Importantly, many mechanisms that induce axonal degeneration do so independently of apoptotic pathways (53), reiterating neurite outgrowth or degeneration as an important, independent evaluation of neural network ‘health’, as demonstrated here.

To measure astrocyte reactivity, we measured immunolabeled GFAP intensity in the 3D CTM. We observed a significant increase in immunolabeled GFAP intensity in irradiated CTMs versus those pre-treated with amifostine (at 24 h) and/or untreated controls (at 48 h). In both cases, this suggests that CTM astrocytes shifted to a reactive phenotype following gamma irradiation and that this reactivity was significantly reduced by the use of amifostine as radiation protectant. Astrocytes are known to mediate neuronal stress, and more specifically oxidative stress, through a number of mechanisms including protection against glutamate toxicity by reducing glutamate uptake at the synapse, increasing glutathione expression, and upregulating oxidative stress response genes via the Nrf2 pathway (55). In acute response to stress or injury, astrocytes shift their phenotype into a ‘reactive’ state, wherein they become hypertrophic and increase expression of kir4.1 and GLAST, genes which promote the reduction of extracellular glutamate and K+ and inhibit inflammation by suppressing excessive microglia activation; actions that provide neuronal protection in most cases (56). Immunolabeled GFAP intensity, ostensibly proportional to GFAP abundance, is a widely used marker for acute astrocytic stress or injury due to its positive proportional relationship with the astrocyte’s reactivity (57–60). It’s worth noting that *in vivo*, immunolabeled GFAP intensity or quantified RNA/protein abundance may be difficult to interpret given the natural variance of GFAP expression across astrocyte subtypes (61–63). For example, astrocytes of the hypothalamus are known to exhibit greater basal expression of GFAP versus those in the amygdala. However, our CTM uses pooled iPSC-derived astrocytes which presumably do not exhibit different basal expression levels across identically prepared and treated samples.

To measure phenotypic activity (i.e., electrophysiological activity) before and after GR-exposure, we measured EAPs or MUAs from 2D co-cultures and 3D CTMs, respectively. We observed a decrease in EAP MFR rate in 2D co-cultures, suggesting disruption of baseline network function. While we observed measurable LDH in both GR-irradiated and untreated cultures, we did not observe a significant increase in LDH as a function of GR-irradiation. This suggests that decreases in MFR were not simply explained by cell death. Moreover, we observed a significant decrease in NBR at the 24 h timepoint, suggesting synaptic disfunction due to radiation exposure. The presence of ROS has been shown to trigger synaptic plasticity through both direct (e.g., modification of NMDA receptors) and indirect (e.g., microglia activation) mechanisms, both *in vitro* and *in vivo* (64, 65), and maladaptive plasticity is linked to pathologies such as Alzheimer’s disease (66). This proof-of-concept data suggests platform technologies such as MEAs and prospectively high throughput calcium imaging can be used to test fundamental hypotheses related to synaptic disfunction following IR-exposure. We did not, however, observe significant differences in MUA in 3D CTMs following GR-exposure. This may be due to 1) relatively low number of MUA ‘events’ being measured, even under baseline conditions, or 2) the functional interactions between astrocytes – known to serve a ROS-buffering role *in vivo* – are sufficiently different in 3D versus 2D systems. Regardless, future work will incorporate penetrating or integrated 3D electrode arrays to enable single-unit EAP recordings from 3D CTMs – an approach that has been demonstrated successful both in engineered tissue models and organoids (67).

The combination of a neuron and astrocytes is sometimes referred to as the neuroglial unit; a statement of cell type necessity in describing neuronal function (68). However, microglia and oligodendrocytes play necessary roles in cortical network function, maintenance of homeostasis, and response to injury (69, 70). Future CTM iterations should build toward a more complete functional cortical tissue ‘unit’, which would integrate the above cell types. This is not without its challenges, including optimization of common maintenance medium and time-staggered delivery of differentiation cues or the necessity to pre-differentiate before seeding. One of the typical disadvantages of engineered cortical tissue models lack the anatomical organization of the cortex. Importantly, the current fabrication strategy is compatible with multilayering of structures containing different cell types and/or cell ratios during or after the initial manufacturing process. This may not, in itself, lead to intra-layer cellular or functional organization, but individual layers may also incorporate growth cues to encourage directional outgrowth and inter-layer functional network formation. An intrinsic challenge facing all tissue models is that of growth factor/metabolic perfusion/diffusion and maintaining a viable and physiologically relevant oxygen gradient (71, 72). It has been reported that a tissue model thickness of greater than 300 µm will be subject to inner-core necrosis (73), thus reducing long term culture and relevance of organoids (74). Importantly, the current model (approximately 300 µm in thickness) was able to support viable differentiated cell growth for up to 64 days, significantly surpassing the 20-day mark showed for other 3D tissue structures without vasculature (71, 74). To extend this model to multi-layered architectures would likely require incorporation of oxygen-releasing biomaterials, spinning bioreactors orbital shakers, and/or the integration biological or fabricated permeable conduits tantamount to vasculature.

## Conclusion

Here, we report on the development of a physiologically relevant cortical tissue model (CTM) that incorporates the fundamental neuroglial unit and approaches the structural and mechanical features of native cortical tissue. To the best of our knowledge, this is the first engineered 3D CTM that has been evaluated following GR exposure using both standard and innovative biological readouts. Our results indicate that our CTM recapitulates standard measures of IR-induced injury reported *in vivo* (e.g., DNA damage, cytotoxicity), and responds to the effect of radioprotectant countermeasure in agreement with clinical studies. Additionally, our data indicates that phenotypic activity measured using *in vitro* MEAs could support addressing fundamental questions regarding the GR effects on cortical networks and serving as a platform for radioprotectant drug identification and development. Importantly, the CTM’s stability and robustness makes it suitable for a variety of radiation sources and durations/repetitions of exposure, including proton beam and mixed exposures for modeling short-to-long term radiation therapy and/or galactic background radiation exposure.

## Methods

### Cell Culture

Neuroblastoma SH-SY5Y cells (provided by Dr. Elaine Lim at UMass Medical School) were cultured in 2D and 3D using Dulbecco’s Modified Eagle’s Medium (DMEM) with GlutaMax (10567014, Thermo Fischer, MA) supplemented with 10% fetal bovine serum (FBS) (26140079, Thermo-Fischer, MA) and 1% penicillin/streptomycin (15140122, Thermo-Fischer, MA), and incubated at 37 °C in a humidified atmosphere containing 5% CO2. SH-SY5Y cells were differentiated in media consisting of NeuroBasal Media (21103049, Thermo-Fischer, MA) with 2% B27 (17504044, Thermo Fischer, MA), 1% GlutaMax (35050061, Thermo-Fischer, MA), 1% penicillin/streptomycin and 1µM retinoic acid (R2625, Sigma-Aldrich, MO). Every four days, 50% of the media volume was exchanged.

Human iPSC derived glutamatergic neurons (BX-0100, BrainXell, WI) and human iPSC derived spinal astrocytes (BX-0650, BrainXell, WI) were co-cultured in a 1:1 ratio in media containing 48% DMEM/F12 (11320-033, ThermoFischer, MA), 48% Neurobasal, 2% B27, 1% N2 supplement (17502048, Thermo-Fischer, MA), and 1% GlutaMax.

In the case of iPSC derived cells, 2D and 3D co-cultures underwent 50% media exchanges every second day for 26 days before treatment with GR. BrainXell astrocyte seeding supplement was included on the seeding day and BrainXell day 4 supplement was included in the Day 4 media exchange at the recommended concentrations. All media changes from day 1 and onward included supplementation with BDNF 10ng/mL, GDNF 10ng/mL and TGF-β1 1ng/mL.

### Morphological Characterization

For SEM characterization, samples were fixed in 4% paraformaldehyde–0.1 M sodium cacodylate solution overnight at 4 °C, washed with deionized distilled water and dehydrated at 4 °C through sequential immersion in a graded series of increasing ethanol concentrations. Samples were subsequently processed with a critical point dryer (Tousimis SAMDRI-795, USA), sputter coated with Au/Pd (Cressington 108, Cressington Scientific Instruments, UK) and analyzed at 10 kV with a Phenom XL G2 Desktop SEM (Thermo Fisher Scientific, MA USA).

### CTM preparation

Collagen hydrogels were prepared, as previously described (36, 37), based on acid solubilized type I collagen extracted from rat-tail tendon (2.05 mg/ml, First Link Ltd., UK). Dense collagen scaffolds were produced by the plastic compression method (38). Briefly, 3 ml of 10×Dulbecco’s modified Eagle medium (DMEM; Sigma Aldrich, MA) was added to 12 ml of type I collagen solution, which was immediately neutralized by adding 5 μl aliquots of 5M NaOH (Fisher Scientific, MA). Stem Cell Qualified ECM Gel (CC131-5ML, Sigma Aldrich, MA) was then added at a 1:80 (volume/volume) ratio, before transferring the collagen-based solution to 24 well plate wells and incubated at 37 °C for 75 min to polymerize. For cell-seeded constructs, cell suspension was added to neutralized collagen solution. Initial experiments to optimize the experimental design were carried with SH-SY5Y cells, while hiPSC derived SMN with spinal astrocytes were seeded in neutralized collagen solution at a density of 4×105 cells/ml. After incubation, the highly hydrated collagen gels were detached from the mold and placed upon a nylon mesh, supported by a stainless-steel mesh on top of blotting paper to absorb the liquid excess (Fig. 1B). Dense collagen sheet scaffolds were then produced by applying an unconfined compressive stress of 1 kPa for 5 min at room temperature. Following compression, each 24-well plate-sized construct was created using a 6 mm tissue biopsy punch (33-36, Integra, NH), creating 5 samples. Each punched sample was gently transferred to an individual well of a 24 well tissue culture plate and cultured in the described tissue culture medium. For acellular constructs, collagen gels were prepared into a custom-made rectangular mold (43 × 50 mm2) and similarly compressed as described after curing at 37 °C (Ghezzi et al. 2012).

### Mechanical Characterization

Compressive mechanical tests were carried out on cylindrically shaped constructs using a UniVert (CellScale, ON, Canada) instrument equipped with a 10 N load cell. Cylindrical collagen constructs were assessed after preparation. All tests were carried out in displacement control with a cross-head speed of 0.01 mm/s. Measurements were conducted at room temperature while maintaining constant sample hydration using drops of distilled water. The specimens’ diameter (5.13±0.4 mm) and height (5.7±0.27 mm) were measured using a digital caliper. Specimens were tested using two parallel nonporous compression platens at up to 80% strain without any evidence of specimen unfolding (n=5). The stress was calculated by normalizing the recorded force against the initial resistance area of the specimen, and the strain was calculated by normalizing the displacement against the initial height of the specimen. The compressive modulus values were computed from the slope of the linear region (20-50% strain) of the stress-strain outputs, as previously reported (39).

### CTM GR treatment

Irradiation with ^60^Co gamma rays was carried out at the UMass Lowell Radiation Laboratory. Both SH-SY5Y and hiPSC-derived seeded samples at day 26 in culture were transported to the high-dose rate gamma cave radiation facility in a Styrofoam cooler with a 37 °C internal environment. The gamma cave provides a shielded room of approximately 512 ft^3^, where samples were treated with GR in the tissue culture plate at a dose rate of 2.5 Gy/min for an elapsed treatment time of two min at room temperature.

### Analysis of cellular viability

Media samples were retained at 24 h following GR treatment. Cell death was quantified based on the abundance of lactate dehydrogenase (LDH) between treated and non-treated samples (MAK066, Sigma-Aldrich, USA). Absolute LDH content was determined with a standard curve. The assay was run according to the protocol provided by the manufacturer. Absorbance values for both assays were measured using a SpectraMax M2 plate reader (Molecular Devices, PA).

### Immunocytochemistry

At 30 min post treatment 3D samples were fixed for assessment of nuclear damage and at 24 h post treatment 3D samples were fixed for assessment of cell morphology. All samples were fixed with 4% paraformaldehyde for 60 min. After three washes with 1X phosphate buffed saline (PBS) (Thermo-Fisher, MA), all samples were blocked in 1% bovine serum albumin (Sigma-Aldrich, MO) for 60 min and permeabilized with 0.2% Triton X-100 (Sigma-Aldrich, MO) for 30 min before another three washes with PBS.

### DNA damage markers

Samples were incubated with mouse monoclonal anti-γ-H2AX (EP854(2)Y, Abcam, UK) (0.7 µg/mL) and anti-53BP1 (A300-272A, Thermo-fisher, MA) antibodies overnight followed by three washes in PBS, and then with donkey anti-rabbit IgG antibody (A21206, Thermo-Fischer, MA) (2 µg/mL) and Goat anti-mouse IgG1 secondary antibody (A-21126, Thermo-Fischer, MA) for 1 h at room temperature. Following incubation with secondary antibodies samples were washed three times with PBS, stained with DAPI for 20 min, and washed three times with PBS less than 1h prior to imaging. Intensity of the phosphorylated histone γ-H2AX and 53BP1, a maker of DNA double stranded breaks (DSB), was quantified using custom ImageJ (NIH) protocols. Images were acquired using identical microscope settings across samples and adjusted such that no pixels in any image was outside of the dynamic range of the imaging hardware. Briefly, the ImageJ analysis protocol used the nuclear marker DAPI as a mask to identify individual objects, then returned the mean grey scale value for all objects detected with an area between 11 and 300 pixels. Axon number and length (βIII-tubulin), neuron and astrocyte nucleus size and circularity (DAPI and NeuN), as well as relative density of pre/post-synaptic markers compared to untreated CTMs using custom ImageJ (NIH) protocols.

### Cellular morphology markers

Samples were stained with NeuN (MAB377[mouse]/ ABN78 [rab]), EMD Millapore), βIII-tubulin, GFAP (AB4674, Abcam), synaptophysin, and PSD95. After incubation in primary antibody samples were washed and incubated for 1 hr in one of the following corresponding secondary antibodies. Samples were imaged after a final round of washes. 2D and 3D samples were washed with ice cold PBS three times and fixed with 4% paraformaldehyde for 60 min. On the day of staining, the cells were first permeabilized with 0.5% Triton X-100 1 h while nonspecific binding sites were blocked with 4% normal goat serum for 2 h. A solution of primary antibodies against glial fibrillary acidic protein (GFAP:1:250; Abcam, Cambridge, MA, USA), neuronal marker (NeuN:1:250; Abcam, Cambridge, MA, USA) in PBS and 4% NGS was incubated on the cells overnight. The following day samples were washed three times with ice cold PBS for 10 min each. Then samples were incubated for 2 h in a solution of secondary antibodies in PBS with of 4% NGS conjugated to their corresponding species with the following wavelengths: 647, 555, 488, and 405 nm. Samples were imaged with a Leica S8 confocal microscope (Leica Biosystems, Danvers, MA, USA). All images were analyzed with ImageJ (NIH, USA).

### Extracellular electrophysiology recordings

Extracellular electrophysiological recordings were carried out using the Axion Maestro Pro multiwell recording platform (Axion Biosystems, Atlanta, GA). This format was used for both 2D hiPSC co-cultures and 3D CTMs. In both cases, voltage measurements were recorded from all seeded wells simultaneously at a sampling rate of 12.5 kHz and filtered using a 1-pole Butterworth bandpass filter (0.1 Hz-5 kHz). In the case of 2D co-cultures, further digital filtering was applied; a 1-pole Butterworth bandpass filter (250-3000 Hz). To identify EAPs in 2D co-cultures, filtered continuous recordings were subject to an adaptive threshold set to ±5.5 standard deviations of the channel’s adaptive RMS. Mean firing rates (MFR) and average electrode yield (AEY) were calculated using Axion’s NeuralMetric software and MFRs from 10-min recordings were compared to their baseline values through difference-over-sum (DoS) normalization. In the case of 3D CTM recordings, ±4(sigma) thresholds were calculated based on RMS calculated for each electrodes recording in total and threshold crossings were defined as multiunit activity (MUAs). Graphing and statistical tests were carried out using OriginPro (OriginLab Corporation, MA, USA). M Statistical comparisons were made by Mann–Whitney *U* test, with *p* < 0.05 considered significant.

### Statistical Analysis

R Studio packages ggplot2 and ggpubr were used for plotting and statistical analysis. Subsequent to either Shapiro–Wilk or Kolmogorov–Smirnov normality test, a Mann-Whitney U-test was used for unequal variances was used to assign p-values to the OTUs when comparisons between two groups were made in the case of DNA damage assessments, cytotoxicity, and cellular morphology. For comparisons across more than two conditions a Kruskal-Wallis ANOVA test with a post-hoc Dunn’s test of multiple comparisons was used. Values of *p<0.05; **p<0.01; ***p<0.001 and ****p<0.0001 were considered statistically significant.

## Notes

### Competing Interest Statement

The authors have declared no competing interest.

